# Tracing two causative SNPs reveals SARS-CoV-2 transmission in North America population

**DOI:** 10.1101/2020.05.12.092056

**Authors:** Xumin Ou, Zhishuang Yang, Dekang Zhu, Sai Mao, Mingshu Wang, Renyong Jia, Shun Chen, Mafeng Liu, Qiao Yang, Ying Wu, Xinxin Zhao, Shaqiu Zhang, Juan huang, Qun Gao, Yunya Liu, Ling Zhang, Maikel Peopplenbosch, Qiuwei Pan, Anchun Cheng

**Author notes:** These authors contribute equally to this work (Xumin Ou, Zhishuang Yang). Correspondence Author: Anchun Cheng; Institute of Preventive Veterinary Medicine, Sichuan Agricultural University, 611130 Chengdu, China; (A.C.).

## Abstract

During the COVID-19 pandemic, precisely tracing the route of the SARS-CoV-2 transmission in human population remains challenging. Because this RNA virus can mutate massively without a specifically tracing maker. Herein, using a geographic stratified genome-wide association study (GWAS) of 2599 full-genome sequences, we identified that two SNPs (i.e., 1059.C>T and 25563.G>T) of linkage disequilibrium were presented in approximately half of North America SARS-CoV-2 population (p = 2.44 x 10^−212^ and p = 2.98 x 10^−261^), resulting two missense mutations (i.e., Thr 265 Ile and Gln 57 His) in ORF1ab and ORF3a, respectively. Interestingly, these two SNPs exclusively occurred in the North America dominated clade 1, accumulated during mid to late March, 2020. We did not find any of these two SNPs by retrospectively tracing the two SNPs in bat and pangolin related SARS-CoV-2 and human SARS-CoV-2 from the first epicenter Wuhan or other regions of China mainland. This suggested that the SARS-CoV-2 population of Chinese mainland were different from the prevalent strains of North America. Time-dependently, we found that these two SNPs first occurred in Europe SARS-CoV-2 (26-Feb-2020) which was 3 days early than the occurring date of North America isolates and 17 days early for Asia isolates (Taiwan China dominated). Collectively, this population genetic analysis highlights a well-confidential transmission route of the North America isolates and the two SNPs we newly identified are possibly novel diagnosable or druggable targets for surveillance and treatment.

---

Since human-to-human transmission of severe acute respiratory syndrome coronavirus 2 (SARS-CoV-2), causative agent of COVID-19, was confirmed on 14-Jan-2020(*1*). At 11-Mar-2020, the world health organization (WHO) had officially started to alarm the human COVID-19 pandemic. Recently, over 4 million of human cases have been confirmed globally, in which the most cases were from North America and Europe. Thus, it is very urgent to precisely trace its transmission route that possibly helps public health surveillance (*2*). However, such a specific surveillance marker is infeasible. Because the SARS-CoV-2 mutates massively without a faithful identity to trace that hampers the precise tracing of its transmission route(*3, 4*). Due to the advancement of genome-wide association study (GWAS) to address the complexity of population genetics, a geographic stratified GWAS study was performed using 2599 full-genome sequences of global SARS-CoV-2 population(*5*). We primarily found that two SNPs (i.e., 1059.C>T and 25563.G>T) of complete linkage disequilibrium were presented in approximately half of the North America SARS-CoV-2 strains. Using the two SNPs for a retrospective tracing study, a well-confidential transmission route of the North America isolates was confirmed. The two SNPs we newly identified are possibly novel diagnosable or druggable targets for surveillance and treatment.

## Phylogenetic tree of global SARS-CoV-2

To identify the phylogeographic relationship of global SARS-CoV-2 population (n=2599), a total of 13 clades was confirmed though strain mixture of geographic distribution exists to some extent that was probably caused by international travel. We found that the North America isolates dominated the clade1 (418/521, 80.23%) and three small clades including clade 2, 3 and 7(**Fig. 1**), in which the clade 7 was also evolutionally closed to the bat (RaTG13) and pangolin related SARS-CoV-2. Besides, the clade 4, 5, 6 and clade 13 were dominated by Europe isolates and clade 9, clade 11 and clade 12 were dominated by Asia isolates (**Fig. 1**). Importantly, the phylogenetic tree is inferred by the massively producing mutations, identification of key mutations can possibly provide clues for tracing the transmission route of SARS-CoV-2 (*6*).

**Figure 1.**
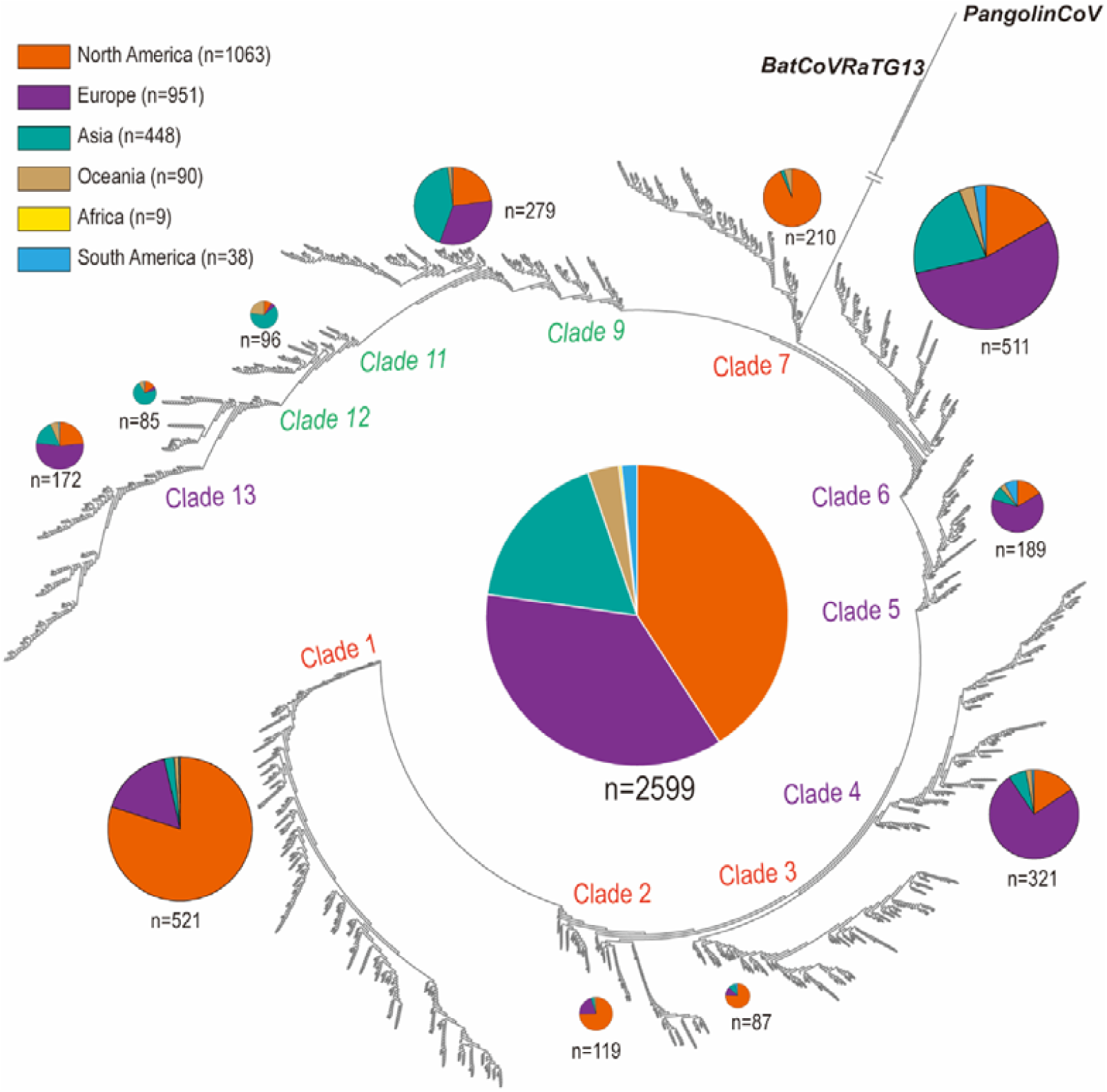
Phylogenetic tree of global SARS-CoV-2. The 2599 full genome sequences (2597 human viruses and 2 wildlife viruses (1 bat and 1 pangolin)) were used. The phylogenetic tree was constructed by Maximum likelihood (ML) method using FastTreeMP software (version 2.1.3, parameter: -nt -gtr -gamma). A total of 13 clades was confirmed though strain mixture of geographic distribution exists to some extent. Specifically, the North America isolates dominated the clade1 (418/521, 80.23%) and three small clades including clade 7, 2 and 3, in which the clade 7 was also evolutionally closed to the bat (RaTG13) and pangolin coronaviruses. The clade 4-6 and clade 13 were dominated by Europe isolates and clade 9, clade 11 and clade 12 were dominated by Asia isolates. The geographic distribution of viruses in each clade were displayed by a pie chart.

## Geographic GWAS study reveals two SNPs of linkage disequilibrium associating the North America SARS-CoV-2 population

Calling key SNPs from the massive mutations apparently requires a GWAS study that has been learned from human GWAS study (also refer to microbial GWAS) (*5*). Because of the mixture of the phylogenetic tree, thus it may be unconfident to perform a phylogenetic stratified GWAS study. This alternatively promoted a geographic stratified GWAS study as the geographic location of individual strain is confident though international travel exists. The mutation features between continents may possibly reflect the incidence of viral mutation emergence in different human hosts(*6*). Using the geographic stratified GWAS study comparing the North America isolates and the outside isolates, we founded that 21 out of 5312 SNPs or INDELs were significantly associated using the Wuhan isolate (GASID: EPI_ISL_402125) as reference genome for SNP calling (Threshold p-value = 1.00 x 10^−15^) (**Fig 2A**). Specifically, the top two SNPs (i.e., 1059.C>T and 25563.G>T) are significantly presented in approximately half of the North America SARS-CoV-2 population (p-value =2.44 x10^−212^ and p=2.98 x10^−261^) and all occur in the clade 1 of North America SARS-CoV-2 population (**Table 1**). Interestingly, the two SNPs are perfect linkage disequilibrium, suggesting the two SNPs concurrently occur in the North America dominating SARS-CoV-2 population (**Fig 2B**). Functional analysis indicted that the two SNPs resulted two missense mutations (i.e., Thr 265 Ile and Gln 57 His) in ORF1ab and ORF3a, respectively. Among these 21 SNPs, we also identified the previously reported 8782.C>T and 28144.T>C SNPs that was consistent to our analysis but had a comparatively low significance (p-value = 4.03 x10^−28^ and 9.73 x10^−33^), resulting a synonymous mutation and a missense mutation (Leu 84 Ser) (*3*) (Table 1).

**Table 1.** List of top 21 hits of SNPs.

| # | SNP | Gene | Effects | P-value | SNP Frequencies |  |  |  | SNP Percentage |  |  |
| --- | --- | --- | --- | --- | --- | --- | --- | --- | --- | --- | --- |
|  |  |  |  |  | North America | North America* | Europe | Asia | North America | Europe | Asia |
| 1 | 25563.G>T | ORF3a | Missense(Gln57His) | 2.98E-261 | 574/1063 | 418/418 | 119/951 | 22/448 | 0.54 | 0.13 | 0.05 |
| 2 | 1059.C>T | ORF1ab | Missense(Thr265Ile) | 2.44E-212 | 479/1063 | 418/418 | 94/951 | 19/448 | 0.45 | 0.10 | 0.04 |
| 3 | 17858.A>G | ORF1ab | Missense(Met5865Val) | 3.57E-117 | 194/1063 | 0/418 | 0/951 | 0/448 | 0.18 | 0.00 | 0.00 |
| 4 | 17747.C>T | ORF1ab | Synonymous | 1.43E-116 | 193/1063 | 0/418 | 0/951 | 0/448 | 0.18 | 0.00 | 0.00 |
| 5 | 18060.C>T | ORF1ab | Missense(Ser5932Phe) | 2.42E-113 | 196/1063 | 0/418 | 0/951 | 4/448 | 0.18 | 0.00 | 0.01 |
| 6 | 29553.G>A | ORF10 | Upstream | 7.26E-51 | 80/1063 | 80/418 | 0/951 | 0/448 | 0.08 | 0.00 | 0.00 |
| 7 | 28882.G>A | N | Synonymous | 6.15E-39 | 53/1063 | 0/418 | 207/951 | 30/448 | 0.05 | 0.22 | 0.07 |
| 8 | 28883.G>C | N | Missense(Gly204Arg) | 6.15E-39 | 53/1063 | 0/418 | 207/951 | 30/448 | 0.05 | 0.22 | 0.07 |
| 9 | 28881.G>A | N | Missense(Arg203Lys) | 2.77E-38 | 54/1063 | 1/418 | 207/951 | 30/448 | 0.05 | 0.22 | 0.07 |
| 10 | 28144.T>C | ORF8 | Missense(Leu84Ser) | 9.73E-33 | 245/1063 | 0/418 | 49/951 | 87/448 | 0.23 | 0.05 | 0.19 |
| 11 | 27964.C>T | ORF8 | Missense(Ser24Leu) | 2.74E-29 | 47/1063 | 47/418 | 0/951 | 0/448 | 0.04 | 0.00 | 0.00 |
| 12 | 11916.C>T | ORF1ab | Missense(Ser3884Leu) | 4.87E-29 | 51/1063 | 0/418 | 0/951 | 2/448 | 0.05 | 0.00 | 0.00 |
| 13 | 8782.C>T | ORF1ab | Synonymous | 4.03E-28 | 245/1063 | 0/418 | 53/951 | 96/448 | 0.23 | 0.06 | 0.21 |
| 14 | 15324.C>T | ORF1ab | Missense(Thr5020Ile) | 5.83E-27 | 2/1063 | 1/418 | 80/951 | 9/448 | 0.00 | 0.08 | 0.02 |
| 15 | 11083.G>T | ORF1ab | Missense(Leu3606Phe) | 3.99E-25 | 71/1063 | 4/418 | 92/951 | 130/448 | 0.07 | 0.10 | 0.29 |
| 16 | 18998.C>T | ORF1ab | Missense(His6245Tyr) | 1.10E-22 | 41/1063 | 0/418 | 0/951 | 2/448 | 0.04 | 0.00 | 0.00 |
| 17 | 29540.G>A | ORF10 | Upstream | 1.10E-22 | 41/1063 | 0/418 | 0/951 | 2/448 | 0.04 | 0.00 | 0.00 |
| 18 | 18877.C>T | ORF1ab | Synonymous(His6245Tyr) | 2.31E-19 | 65/1063 | 0/418 | 10/951 | 8/448 | 0.06 | 0.01 | 0.02 |
| 19 | 29711.G>T | 5'UTR | Downstream | 4.30E-19 | 31/1063 | 1/418 | 0/951 | 0/448 | 0.03 | 0.00 | 0.00 |
| 20 | 1604.AATG>A | ORF1ab | Deletion(delTGA) | 2.87E-17 | 4/1063 | 0/418 | 69/951 | 1/448 | 0.00 | 0.07 | 0.00 |
| 21 | 27046.C>T | M | Missense(Thr175Met) | 3.48E-17 | 2/1063 | 0/418 | 60/951 | 2/448 | 0.00 | 0.06 | 0.00 |

North America\*, clade 1 of the North America SARS-CoV-2 population

**Figure 2.**
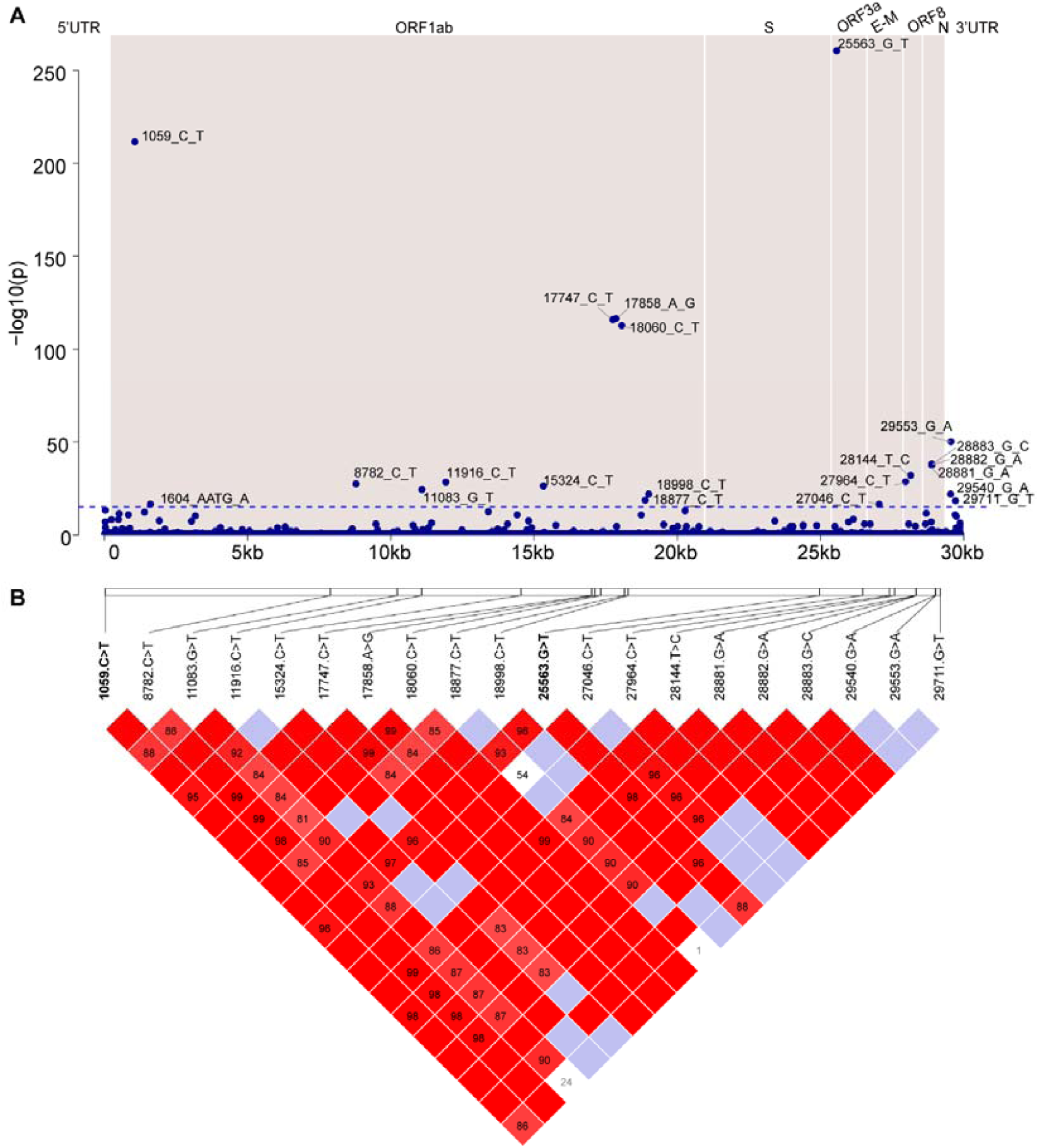
GWAS and Linkage disequilibrium (LD) analysis. **(A)** Manhattan plot comparing the North America SARS-CoV-2 population to other continent isolates. Genomic coordinates are displayed along the X-axis, and -log_10_ of the association p-value for each SNP are displayed on the Y-axis (1.00 x10^−15^). Different blocks indicate different protein encoding region. (**B**) Linkage disequilibrium between SNPs in the SARS-CoV-2. LD plot of any two SNP pairs among the 21 sites (see table 1). The number near slashes shows the genomic coordinates. Color in the square is given by standard (D’/LOD), and the number in square is r^2^ value.

## Retrospective tracing study reveals a well-confidential transmission route of the North American SARS-CoV-2 population

In the North America SARS-CoV-2 population, 54% strains have these two SNPs and in particular 100% for the clade 1 (**Table 1**). Because of the high occurrence of the two SNPs, tracing the two SNPs may provide a well-confidential transmission route of the SARS-CoV-2 in major North America human population. We thus did a retrospective tracing study in our high confidential database (2599 flittered strains) to identify the time sequential of occurring the two SNPs among continents. We found that the first occurring incidence started in France, Europe (26-Feb-2020) which was 3 days early than the occurring date of the North America isolates (29-Feb-2020) and 17 days early for the Asia isolates (Taiwan but not Chinese mainland) (13-Mar-20202, Taiwan, China) (Fig 3) (**Table 2**) (**Fig 3A-G**). Further tracing the accumulating frequencies per day of the two SNPs among continents, we found that during mid to late March, the North America isolates highly accumulated such two SNPs. Besides, the mean number of all accumulating SNPs before or after mid to late March were significantly lower than that during the same period (**Sup Fig S1**). These evidences indicated that the two SNPs were strongly selected in North America SARS-CoV-2 population during mid to late March. The accumulation of the two SNPs may possibly explain the sharp increase of confirmed cases in the North America before early April (data not shown). Because the viruses acquired such two strong selected sites at the same period that may enhance it spread in the North America human population.

**Figure 3.**
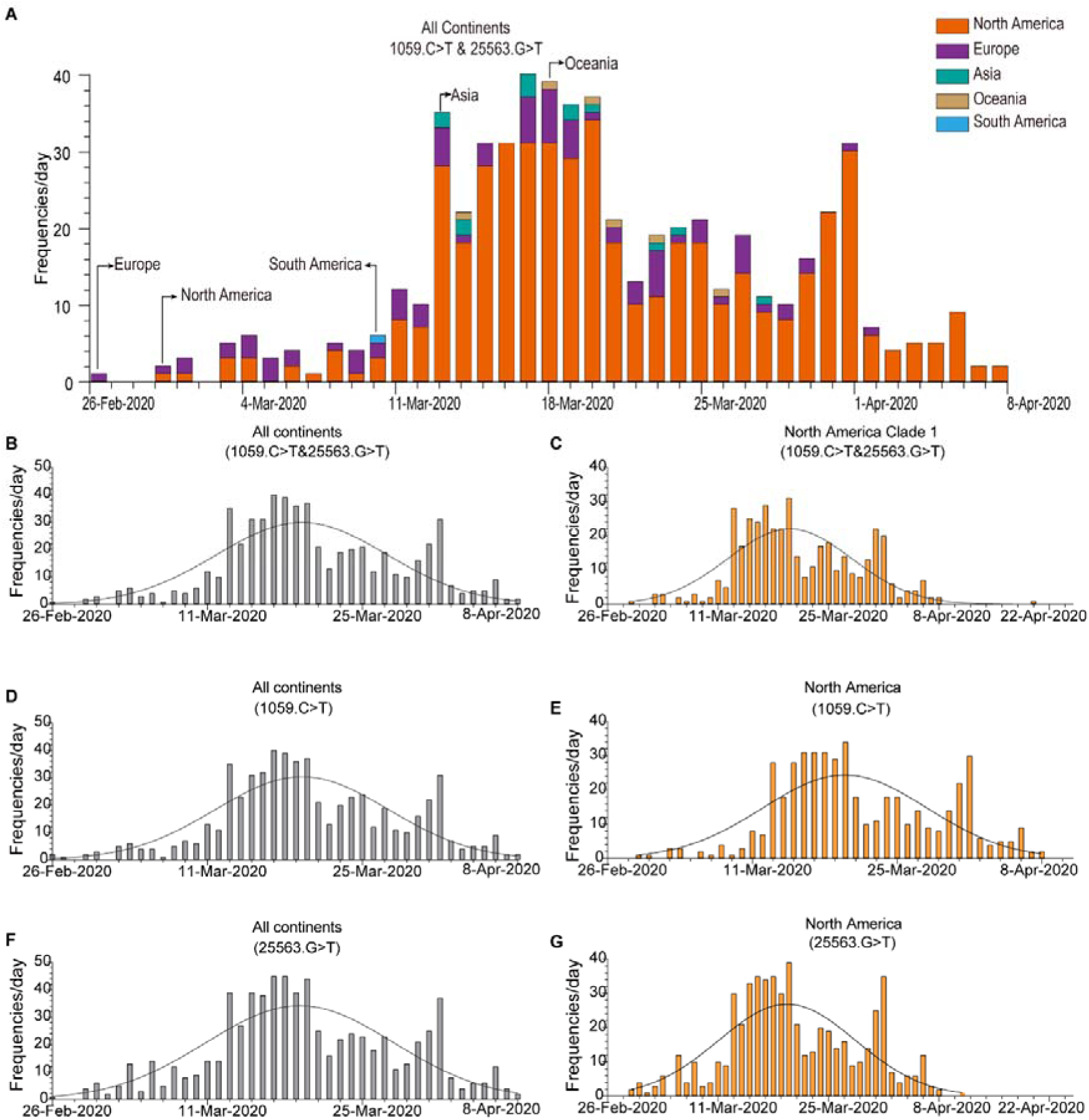
Retrospectively tracing the two SNPs of SARS-CoV-2 between all continents and the North America. (**A**) The time-dependent accumulating plot for frequencies of the two SNPs between continents. The continents are labeled by different colors and the date of first occurrence of the two SNPs are also indicated. (**B-G**) The time-dependent accumulating plot (1059. C>T & 25563. G>T) in either all continents’ SARS-CoV-2 or the North America isolates. Panel **B-C** for occurrence of the two SNPs; panel **D-E** for SNP (1059. C>T) and panel **F-G** for SNP (25563. G>T). Of note, the two SNPs largely accumulated during mid to late march and the most for North America SARS-CoV-2.

Because the SARS-CoV-2 was believably transmitted from wildlife to human before happening of human-to human transmission (*7, 8*) (*3*). We then carefully scrutinized if bat or pangolin coronaviruses had the two mutations before they jumped to human species. But, we did not find any of these two SNPs in the bat or pangolin related coronaviruses because they were highly skeptical for human COVID-19 pandemic ascribing to high sequence similarity (**Sup Fig S2**). Alternatively, in the bat or pangolin coronaviruses, the 1059 site has no or a C>A mutation and the 25563 site has a G>A mutation instead (**Sup Fig S2**). These evidences indicated that the bat and pangolin coronaviruses might undergo a strong selection after jumping to human population at least in the major North America human population.

## CONCLUSION REMARK

Using the two SNPs identified by the geographic stratified GWAS study, we have now reconstructed the well-confidential transmission route of North America SARS-CoV-2 population, that the first transmission incidence with the two SNPs happened at Europe (26-Feb-2020), the North America (29-Feb-2020), then South American (10-Mar-2020), and later Asia (13-Mar-2020) and Oceania(14-Mar-2020). Regardless of the importance of the transmission route, the two SNPs also pose great possibility for epidemic surveillance in North America human population as its high prevalence in the same population. To precisely tracing of the COVID-19 pandemic, development of a differential diagnosis is urgently needed, such as the two SNPs based real-time reverse transcription polymerase chain reaction (PCR) method (*9*).

Moreover, the two SNPs are responsible for two missense mutations. The missense mutations may change the protein function to some extent through we do not provide the evidences but it is generally believed. More mechanical investigations of the functional impact caused by these two SNPs would be enhanced in this aspect as the they are possibly druggable targets. In the future, it is hoped, by detecting to the two SNPs harboring SARS-CoV-2 we estimated, this would help public health surveillance and investigation of their functional impacts would promote novel drug development.

## Materials and methods

### 2.1. Data acquisition

In this study, whole-genome sequences of global SARS-CoV-2 collected from 12-Dec-2019 to 24-Apr-2020 (8:00 GMT + 8) were archived from the database of GISAID Initiative EpiCoV platform (GISAID; https://www.epicov.org). A total 8480 sequences were archived and primarily filtered by the criteria, such as high coverage only (> 29,000 bp), exclusion of low coverage and the sequence with unknown bases (N) inside. The identically redundant sequences was further removed by CD-HIT software (version 4.8.1, parameters: -aL 1 -aS 1 -c 1 -s 1) (*10*). A final 2599 sequences with high confidence were used in this study (**Table S1**).

### 2.2. Phylogenetic analysis

The 2599 full-genome sequences were aligned by MAFFT software (version 7.407, parameter: --auto)(*11*). The phylogenetic tree of the globally 2599 SARS-CoV-2 was constructed by FastTreeMP software (version 2.1.3, parameter: -nt -gtr -gamma) using approximately maximum likelihood (ML) method. A total 13 clades were identified and the geographic distribution of each clade were displayed by a pie chart.

### 2.3 SNP calling

The single nucleotide polymorphisms (SNPs) and small insertion-deletion (INDELs) polymorphisms were detected by MUMmer software (version 3.0, nucmer, show-snps) (*12*) using the Wuhan-Hu-1 strain (GISAID: EPI_ISL_402125, Genbank: NC_045512.2) as a reference genome. To validate identity of the above polymorphisms, raw reads (40 out of 2599 strains, NCBI SRA database) were analyzed by bwa (version 0.7.16a) (*13*) and mpileup program of samtools software (version 1.10)(*14*). The validation was consistently to the polymorphisms detected by the MUMmer software (data not shown).

### 2.4 Genome-wide association study (GWAS) and Linkage disequilibrium(LD) analysis

In order to identify causative SNPs in population of North America SARS-CoV-2 (cases=1063, controls=1536), a geographic stratified genome-wide association study against 5312 mutations was performed using PLINK software (version 1.90) (*15*). The empirical threshold of p-value was suggested to be 9.41 x10^−6^ (0.05/5312=9.41 x10^−6^) calculated by Benjamini & Hochberg method (1995) (*16*), but we further increased the threshold of p-value to 1.00 x10^−15^ for detection of the most causative SNPs. Consequently, 21 significant SNPs were detected (Table 1) and the LD of paring SNPs were estimated and visualized by Haploview software (version 4.1) (*17*).

### 2.5. SNPs accumulating analysis

To analyze the trend of SNPs accumulation during March, 2020, the frequencies of average SNPs accumulation per day was counted. The same counting of the two SNPs (i.e., 1059.C>T and 25563.G>T) in North America SARS-CoV-2 and the other continents were traced by the date of occurring the two SNPs. These analyses were performed by Microsoft® Excel 2016 (**Table S2**).

### 2.6. Statistics

Data from 2.5 were plotted by Graphpad (Version 8.2.1 for Windows, San Diego, CA). The mean differences of the average SNPs accumulation per day during continuous ten-days were analyzed by Mann-Whitney U test (R version 3.6.2). A p-value less than 0.05 is considered as statistically significant.

## Acknowledgements

We thank members of the Institute of Preventive Veterinary Medicine for their valuable suggestion and discussion. **Findings:** This work was supported by grants from China Agricultural Research System (CARS-42-17), Sichuan Veterinary Medicine and Drug Innovation Group of China Agricultural Research System (SCCXTD-2020-18), Science and Technology Program of Sichuan Province (20YYJC4428) and Integration and Demonstration of Key Technologies for Goose Industrial Chain in Sichuan Province (2018NZ0005). **Competing interests:** The authors declare no competing interests. **Data and materials availability:** All data is available in the main text or the supplementary materials.

---

**Sup Figure S1.**
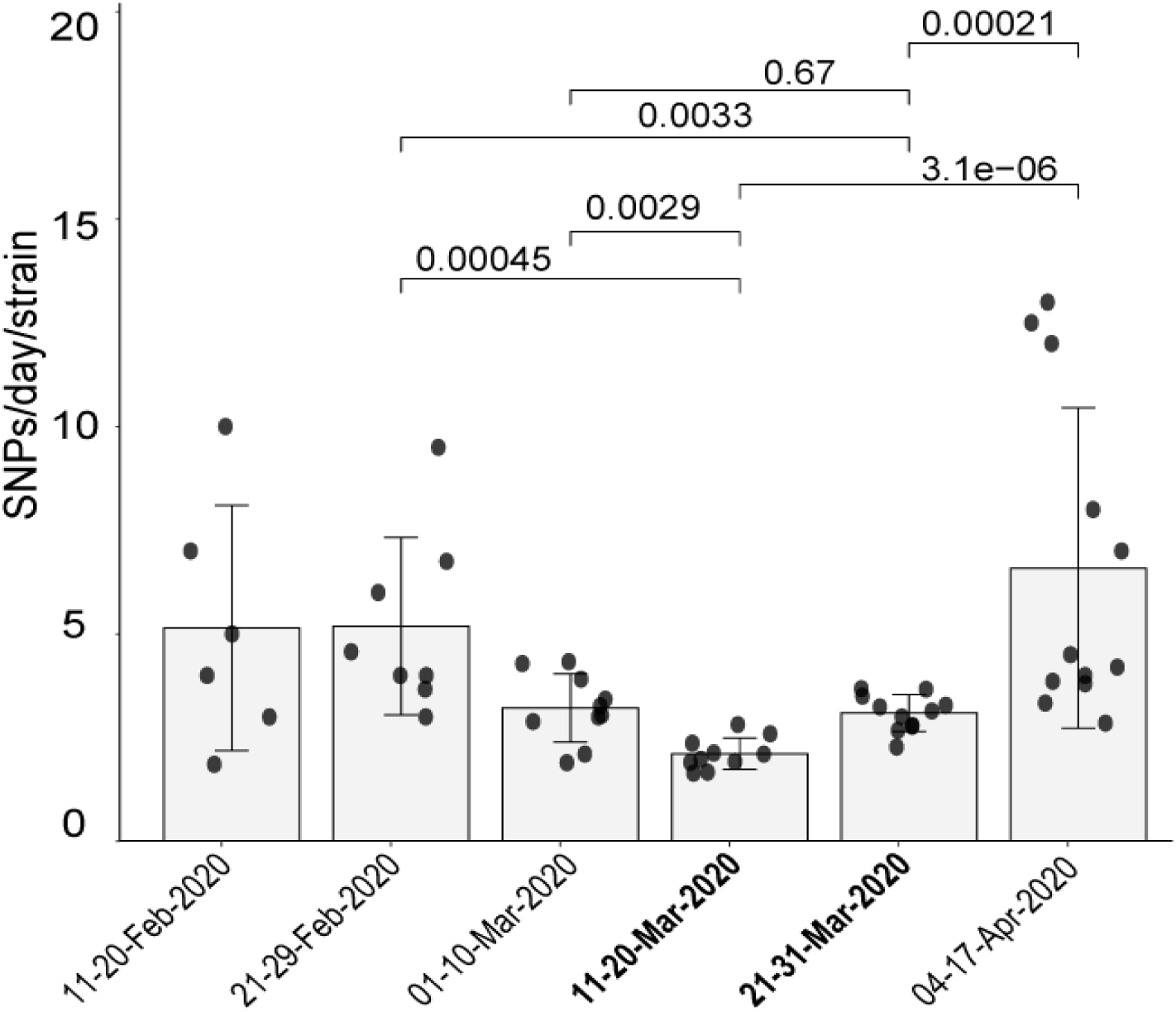
SNPs accumulation during mid to late March, 2020. The mean differences of the average SNPs accumulation per day during continuous ten-days were analyzed by Mann-Whitney U test. A p-value less than 0.05 is considered as statistically significant.

**Sup Figure S2.**
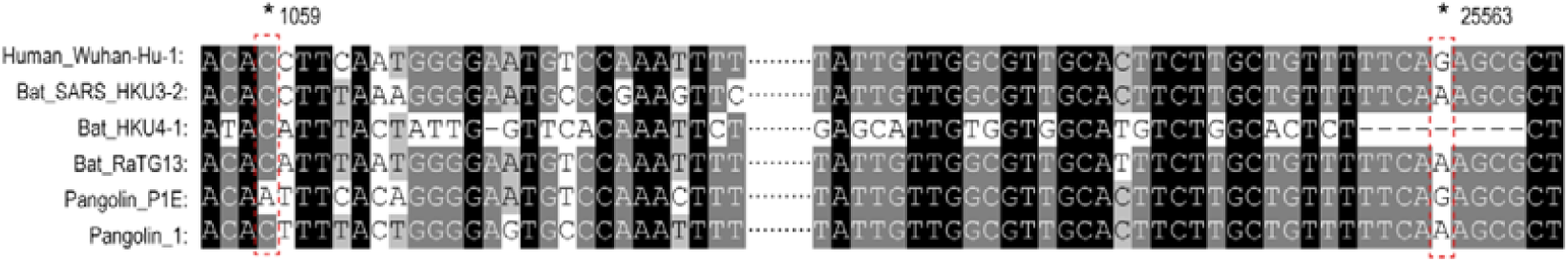
SNP calling in bat and pangolin related SARS-CoV-2 at 1059 and 25563 sites. Sequence alignment between bat and pangolin related SARS-CoV-2 was displayed. Human Wuhan-hu-1 strain was used as reference genome. Strains from bat and pangolin were named with its host name. Differently, at 1059 site, there is only one SNP for pangolin P1E strain (C>A) but not C>T. And the 25563 site has another type of SNP (G>A) instead of G>T.

